# Laws of concatenated perception: Vision goes for novelty, Decisions for perseverance

**DOI:** 10.1101/229187

**Authors:** D. Pascucci, G. Mancuso, E. Santandrea, C. Della Libera, G. Plomp, L. Chelazzi

## Abstract

Every instant of perception depends on a cascade of brain processes calibrated to the history of sensory and decisional events. In the present work, we show that human visual perception is constantly shaped by two contrasting forces, exerted by sensory adaptation and past decisions. In a series of experiments, we used multilevel modelling and cross-validation approaches to investigate the impact of previous stimuli and responses on current errors in adjustment tasks. Our results revealed that each perceptual report is permeated by opposite biases from a hierarchy of serially dependent processes: low-level adaptation repels perception *away* from previous stimuli; high-level, decisional traces attract perceptual reports *toward* previous responses. Contrary to recent claims, we demonstrated that positive serial dependence does not result from *continuity fields* operating at the level of early visual processing, but arises from the inertia of decisional templates. This finding is consistent with a Two-process model of serial dependence in which the persistence of read-out weights in a decision unit compensates for sensory adaptation, leading to attractive biases in sequential responses. We propose the first unified account of serial dependence in which functionally distinct mechanisms, operating at different stages, promote the differentiation and integration of visual information over time.

## Introduction

From moment to moment, human perception is not a high-fidelity replica of sensory input but relies heavily on past experience and short-term dependencies: What happened a moment ago has a strong impact on how we perceive the present.

Vision, for example, is inherently contaminated by the immediate past: low-level visual features such as brightness, color, or motion are all perceived depending on their temporal context ^1–3^, and when the stimulation history changes, identical stimuli may well lead to dissimilar percepts ^1,4,5^ Such short-term dependencies are not peculiar to visual perception, but permeate a wide range of cognitive processes, including attention ^6–8^, decision-making ^9–11^, memory ^12,13^, confidence in performance ^14^ and motor behavior ^15^. This implies that, at multiple stages, our cognitive system is anchored and calibrated to the recent history of sensory and decisional processes.

From a phenomenological perspective, short-term dependencies may carry opposite effects on perception: Negative (repulsive) effects are observed when perception is repelled away from previous stimuli ^1,4,16,17^ Positive (attractive) effects, instead, arise when consecutive stimuli appear more similar than they really are and, thus, past representations persist and “attract” current perceptions. A recent collection of studies suggests that the latter, positive form of serial dependence dominates visual perception ^18^.

The notion of positive serial dependence in visual perception comes from evidence of systematic errors in adjustment tasks pulled toward stimuli seen one or a few trials before ^18,19^ This phenomenon has been observed for features encoded at multiple stages along the visual hierarchy, including for basic stimulus features, such as orientation,^19^ spatial position ^20^ and numerosity,^21^ but also for more complex stimuli, such as faces,^22^ emotion ^23^and facial attractiveness.^24^ Such ubiquity of positive biases has led some authors to hypothesize that serial dependence is an intrinsic mechanism through which our visual system exploits temporal correlations and contextual redundancies by merging similar stimuli, slightly changing over time, into a coherent and “continuous” perceptual field ^18,19^.

Although the idea of a continuity field fits neatly with recent findings, it stands in sharp contrast with other, widely documented forms of negative dependence. Examples of negative serial dependence are well-known phenomena of visual adaptation and aftereffects ^17^. In visual aftereffects, previous exposure to persistent features, such as orientation ^4^, color ^17^ or motion ^1^, causes subsequent stimuli to appear as shifted away from the exposed feature: a stationary object will seem to be moving downward after exposure to upward motion; a vertical grating will look as if it is tilted clockwise after exposure to anticlockwise orientations. These negative biases go plainly in the opposite direction to that predicted by the continuity field and thus the question arises as to how these two contrasting forces, one attractive and the other repulsive, may coexist in perceptual processing ^25^.

While there is a wealth of evidence linking negative aftereffects to transitory alterations in the response properties of relevant populations of sensory neurons ^26–29^, less is known about the mechanisms regulating positive serial dependence: Is there a shared mechanism responsible for both types of dependencies or are the two phenomena related to distinct and independent stages of visual processing ^30^?

It has been recently proposed that negative and positive history biases originate at different stages of the processing stream ^30^ According to this view, positive serial dependence would result from post-perceptual biases in a working memory buffer where information about recent stimuli is maintained. However, this account remains mostly theoretical and the exact role and nature of the post-perceptual processes involved has yet to be established ^30^.

In the present work, we provide compelling evidence for the existence of two, separate and functionally distinct stages of visual serial dependence ^30–32^, determined by the independent history of sensory and decisional processes. Contrary to previous claims, we demonstrate that there is no such thing as the continuity field in low-level visual perception and that attractive biases are not the result of biased memory representations for previous stimuli; instead positive serial dependence derives from residual traces of past decisions. Using the method of adjustment, in a series of experiments we measured the ability of human participants to reproduce the orientation of consecutive stimuli ^19^ Multi-level analyses and cross-validation techniques were adopted to demonstrate that past perceptual decisions, rather than the physical orientations of previously seen stimuli, are the effective sources of response biases. We found that stimulus-driven serial dependence disappeared when participants were explicitly instructed to withhold their response; in fact, under this circumstance, repulsive biases emerged. This result was corroborated by the striking finding that, after streaks of non-reported orientations, positive serial dependence was still dictated by the most recent reported perceptual decision. This dependency on previous decisions was not the mere consequence of hysteresis in motor responses as it was strongly modulated by stimulus-specific features, such as their contrast. We further demonstrated that the effect of decisional traces generalizes to other tasks and stimuli.

In light of our results, we propose a hierarchical model of short-term dependencies where repulsive biases emerge as a result of adaptation of low-level sensory neurons and positive serial dependence arises from the persistence of read-out biases at a higher-level, decisional stage. Whereas rapid negative aftereffects in response to task-irrelevant stimuli may promote perceptual sensitivity to relevant environmental changes, positive decisional traces may reflect the reinstatement of the most informative sensory channels in the recent past, thus increasing the efficiency of decisional templates, as reported in perceptual learning studies ^33^.

Our work provides a new perspective on the history of sensory and decisional processes by reconciling opposite forms of short-term dependencies into a unified account of concatenated perception and decision-making.

## Results

### Experiment 1: Response vs. Stimulus

A primary goal of this study was to determine whether serial dependence in visual perception is induced by the physical stimuli or by the response decision in the previous trial. Recent work has shown that past stimuli systematically bias the decision about the present sensory input ^19,30,34^ However, the history of decisions may itself represent a source of perceptual bias that combines with or even outweighs the impact of previous stimuli ^30–35^,. To test this possibility, in the present experiment (Experiment 1) we compared the ability of sensory stimuli and perceptual decisions to predict future errors in an orientation-adjustment task ^19^.

Ten participants were presented with peripheral Gabor stimuli (50% Michelson contrast), oriented randomly between 0° and 165° off vertical (in steps of 15°), and were asked to reproduce the perceived orientation by adjusting the tilt of a response bar (see Figure 1A). To compare the influence of past stimuli and responses, we built two competing models. In one model, we predicted errors in the adjustment responses with the difference between the previous stimulus orientation and the current stimulus orientation (delta stimulus, Δ*S*). In the alternative model, we predicted errors with the difference between the previous decision and response and the current stimulus orientation (delta response, Δ*R*). In both models, errors were fitted to a derivative of Gaussian function (DoG) in a multilevel modeling framework (see Methods) and the amplitude parameter of the DoG (α) was used to quantify the magnitude of serial dependence and to test its significance.

**Figure 1.**
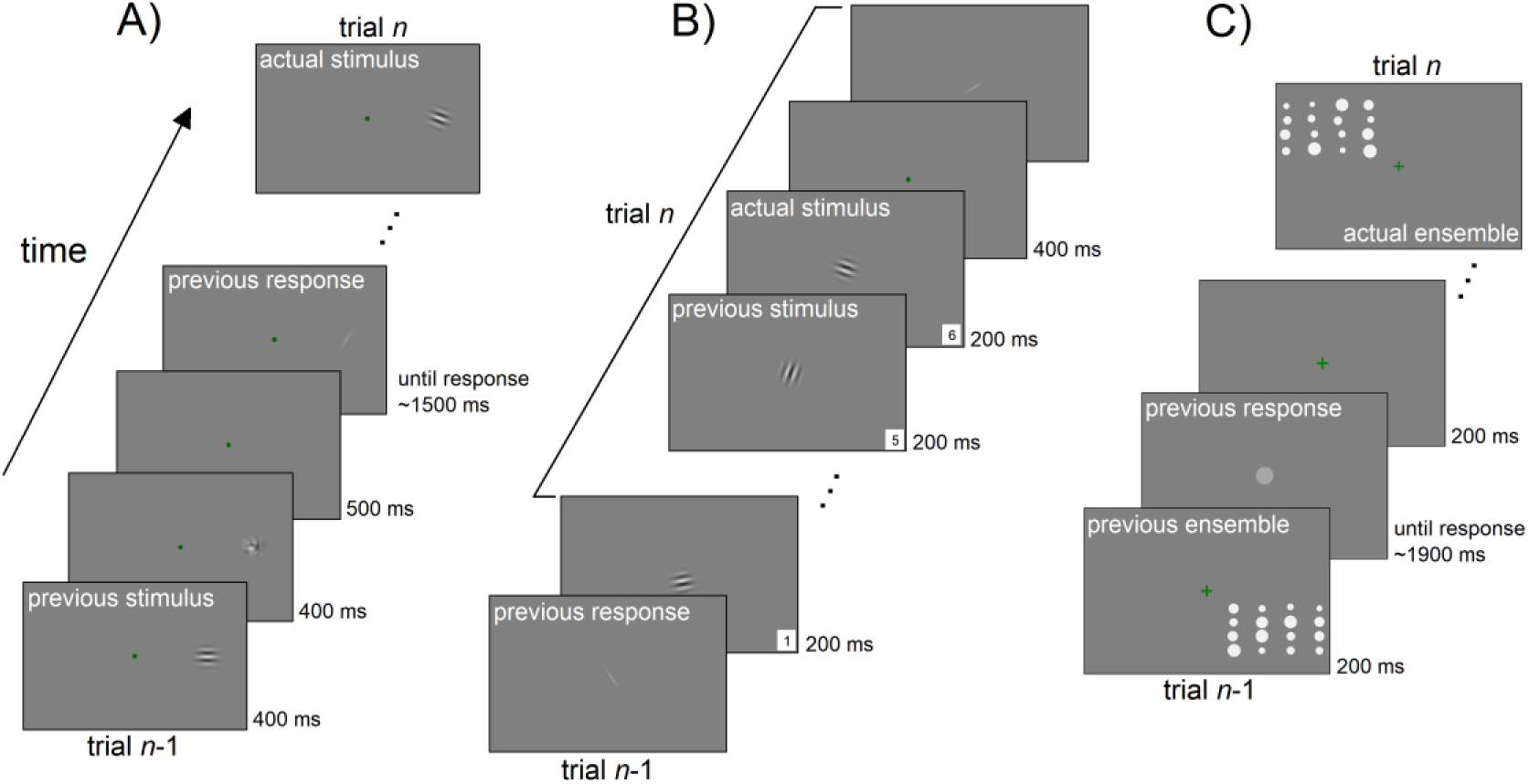
Sequence of events in the different experiments. A) An example of a standard trial in Experiment 1-3,5 and 6. Participants viewed peripheral Gabor stimuli and reported their orientation by adjusting a response bar. In Experiment 2, stimuli were presented at the fovea. In Experiment 1, 3 and 6, stimuli were presented to either the left or right on separate runs. In Experiment 5, the two positions were intermixed randomly. B) A standard sequence of stimuli in Experiment 4. In this experiment, responses were interleaved with sequences of new stimuli. In a single trial, participants had to pay attention to an entire sequence of stimuli and to report the orientation of the last one. The sequence ended after the 6^th^ stimulus in 80% of the trials and stopped earlier (at a random position in time) in the remaining 20% (catch trials). C) Example of one trial in the ensemble coding task of Experiment 7. Participants viewed an ensemble of 16 dots in one of the four quadrants and had to report the average size by adjusting the size of a response dot.

Serial dependence for the stimulus model Δ*S* was small and marginally significant (α = 1.19°, *p* = 0.06) and had a peak at 18.5° (Figure 2A, left panel), in keeping with previous reports ^30^. Importantly, when errors were conditioned on previous responses (Δ*R*), the magnitude of serial dependence was about a factor of two larger than the one observed for Δ*S* (α = 2.31°, *p* < 0.01, peak at 21.8°, see Figure 2A, right panel).

**Figure 2.**
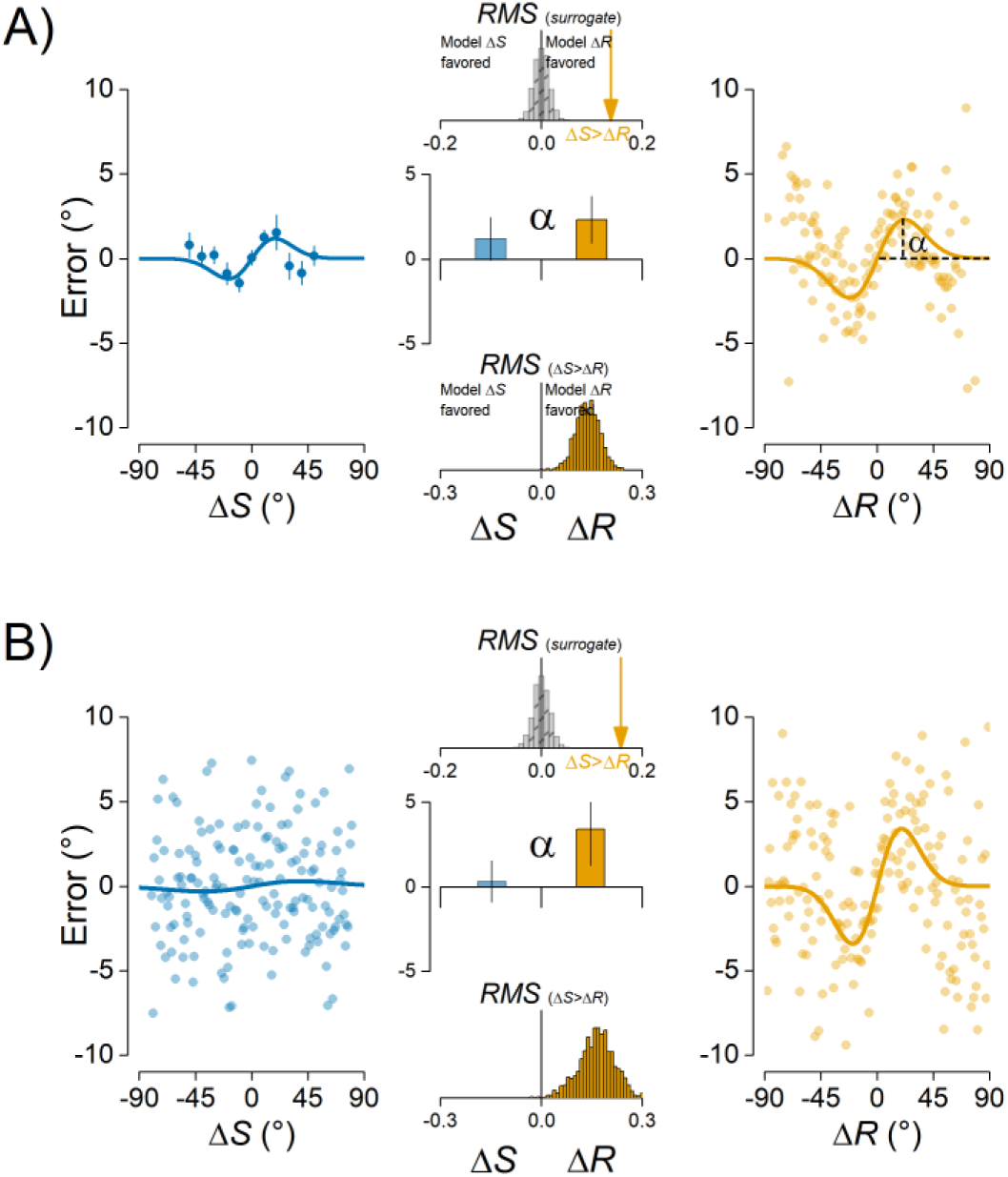
Past responses predict perceptual errors better than previous stimuli. A) Errors conditioned on previous stimuli (Δ*S*, left panel, blue dots and line) and responses (Δ*R*, right panel, orange dots and line) in Experiment 1. The magnitude of serial dependence (α, central bar plot, middle panel) was higher for ΔR compared to ΔS. The ΔR model was also superior in terms of the overall fit (top histogram, middle panel), as shown by the significantly higher *rms* for the Δ*S* model (Δ*S*>Δ*R*, orange arrow) compared to a null distribution obtained by shuffling previous stimuli and responses (Monte Carlo statistic, grey bars). In addition, the Δ*R* had a significantly higher ability to predict new data, as shown by the distribution of *rms* differences (Δ*S*>Δ*R*) obtained in the test phase of the cross-validation statistics (orange bars, bottom histogram). B) Results of Experiment 2 at the fovea. Same representation as in panel A). Lines are best fitting DoG curves from multilevel nonlinear modeling. Bars in bar plots are 95% confidence intervals on the parameter estimates.

To evaluate the best model of serial dependence, we followed two steps: 1) we assessed which model provided a better fit of participants’ errors by computing the difference in the root mean square error (*rms*) between ΔS and Δ*R*, and 2) we compared the ability of each model to predict new data with a cross-validation procedure (see Methods). The model comparison in step 1) favored the Δ*R* model over the ΔS model (*rms*(Δ*S*-Δ*R*) = 0.137, *p* < 0.01, Monte Carlo permutation test, *n* = 1000; Figure 2A, middle panel, top histogram) and in the cross-validation of step 2), the performance of the ΔR model was significantly higher as compared to the Δ*S* model (*p* < 0.05, Figure 2A, middle panel, bottom histogram). Thus, both procedures favored the ΔR model, showing that conditioning on previous responses increased the explanatory and predictive ability of the model.

### Experiment 2: The effect of previous decisions at fovea

The results of Experiment 1 provided strong evidence that previous responses constitute a more efficient predictor of serial dependence than the relative orientation of consecutive stimuli, suggesting that attractive biases may indeed result from the history of perceptual decisions. One possibility is that decisional traces bias perception more than previous stimuli when the sensory input is uncertain.^35^ Because orientation sensitivity degrades as a function of eccentricity ^36,37^, the presentation of Gabors in the periphery of the visual field may have favored decisional biases and lowered the contribution of previous uncertain stimuli. Therefore, it could be hypothesized that when stimulus uncertainty decreases, the physical orientation of previous stimuli becomes a more reliable predictor of perceptual errors compared to the previous decision and response. To address this possibility, we performed a second experiment (Experiment 2) in which the same stimuli of Experiment 1 were presented at the fovea. Although there is evidence of stimulus-induced attractive biases at or near the fovea ^19^, a direct investigation of the impact of previous responses for central stimuli is lacking. In addition, to further reduce stimulus uncertainty (e.g., to increase orientation discriminability) stimuli were presented with higher spatial frequencies (see Methods) ^38^.

Following the procedure of Experiment 1, in six new participants we compared the contribution of ΔS and Δ*R* for centrally presented stimuli. In addition, to rule out any confound due to the comparison of one model - the Δ*S* - built with a discrete predictor against a second model - the Δ*R* - built with a continuous predictor^39^, we varied the relative orientation of consecutive stimuli in the range of ±90° with steps of 1°, making the Δ*S* predictor fully comparable with the Δ*R*.

Contrary to the uncertainty hypothesis, the results of this experiment revealed no serial dependence for Δ*S* at the fovea (α = 0.30°, *p* = 0.623), but a strong dependence of perceptual errors on previous responses (α = 3.40°, *p* α 0.01, peak at 19.5°; Figure 2B). Moreover, model comparison favored the Δ*R* model (*rms*(Δ*S*-Δ*R*) = 0.157, *p* < 0.01; Figure 2B, middle panel, top histogram) which also predicted new data with higher accuracy than the Δ*S* model (*p* < 0.01; Figure 2B, middle panel bottom histogram), thus fully confirming the results of Experiment 1.

The first two experiments yielded two important results: 1) previous perceptual decisions represent a better predictor of serial dependence than previous stimuli and 2) decisional traces affect perception at varying eccentricities whereas the effect of previous stimuli seems to vanish at the fovea.

### Experiment 3: Contrasting forces on serial dependence

Our model comparison revealed that perceptual decisions and behavioral responses are critical, if not fundamental, for serial dependence in orientation discrimination. In their seminal work, Fischer and Whitney ^19^ reported sequential effects of stimulus presentation in the absence of prior motor responses, ruling out the contribution of post-perceptual effects on serial dependence. However, in their experiment, participants were not explicitly instructed to withhold their response and this may have still promoted post-perceptual processing of the stimuli (e.g., in the form of implicit decisions). As a consequence, even in the absence of an actual overt motor response, perceptual decisions might have occurred, leading to lingering effects during the inter-stimulus interval, in turn affecting perception of the subsequent Gabor stimulus. Following this reasoning, we predicted that, if serial dependence emerges from previous decisions, then it should disappear when participants are explicitly instructed to withhold their response.

To test this hypothesis, twelve participants performed an experiment similar to Experiment 1 but including 40% of “catch” trials in which no response had to be made. Crucially, catch trials were explicitly signaled by a black circle appearing at the same location and time as the response bar in the standard trials, indicating that no response - and therefore no perceptual decision - had to be made. Serial dependence was analyzed as a function of Δ*S* after both catch and response trials, and as a function of Δ*R* after response trials only.

After response trials, there was a marginally significant serial dependence for Δ*S* (α = 1.18°, *p* = 0.059, peak at 19.8°) and a strong dependence Δ*R* (α = 3.08°,*p* < 0.001, peak at 14.1°). Interestingly, not only attractive biases were absent after catch trials, but ΔS revealed a repulsive effect on current errors (α = -0.95°, *p* < 0.01, peak at -41.5°) that was significantly different from the effect after response trials (α(catch) vs. α(response), *p* < 0.01; Figure 3), ruling out the possibility that serial dependence for stimuli disappeared merely because of an intervening stimulus (the black circle).

**Figure 3.**
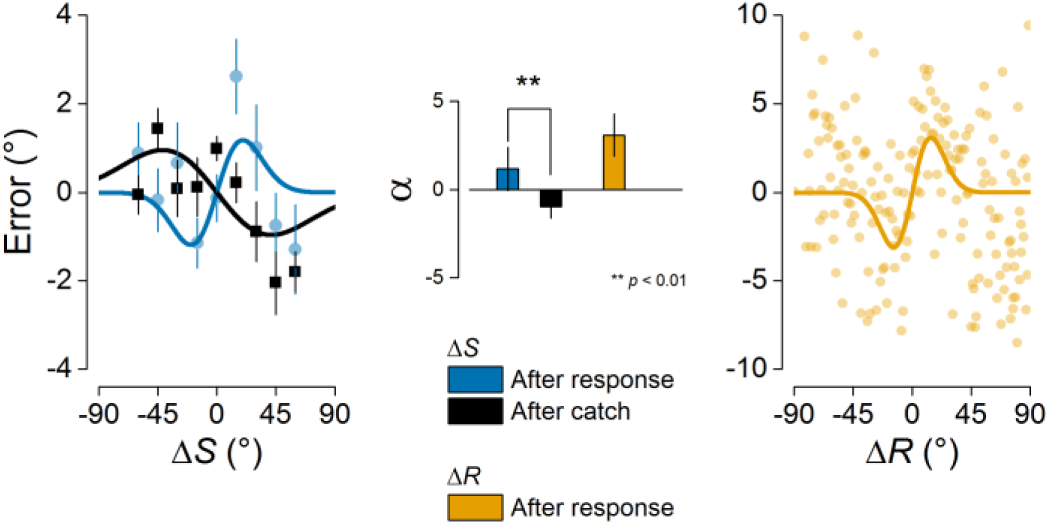
Negative serial dependence after non-response trials. Results of Experiment 3 showing that Δ*S* had an opposite effect after catch trials (40%) or response trials (60%). Positive serial dependence was observed after responses (left panel, blue dots and line), while negative aftereffects emerged after catch trials (left panel, black squares and line). The magnitude of negative and positive dependencies (α) was significantly different (blue vs. black bar, central bar plot). As in Experiment 1, positive carry-over effects were observed when conditioning errors on previous responses (left panel, orange dots and line; middle panel, orange bar).

Confirming our hypothesis, this experiment showed that prior perceptual decisions are fundamental for positive serial dependence and, even more importantly, it revealed that withholding decisions and responses leads to the emergence of repulsive biases, resembling those observed in negative aftereffects ^1,4,28^.

### Experiment 4: Testing a Two-process model of serial dependence

The opposite effect of past stimuli and responses is in sharp contrast with the idea of serial dependence as a low-level visual mechanism ^19^ In line with what would be predicted by the continuity field, serial dependence would originate from changes in the response properties of low-level visual neurons, and therefore it should be immune to post-perceptual processes. However, our results revealed two mechanisms regulating serial dependence, one repulsive, after the mere exposure to sensory stimuli, and the other attractive, observed after perceptual reports. These contrasting forces on visual perception suggest that positive and negative carry-over effects may originate at different stages of the processing stream ^30–32^.

To explore the plausibility of multiple stages of serial dependence, we defined two data-generating models (Figure 4) that implemented serial dependence either as a consequence of low-level processes (Gain model) or as a combination of opposite forces exerted by mechanisms at different stages (Two-process model). In both cases, perception was modeled as an encoding-decoding process performed by a hierarchy of units based on recent models of visual processing ^33,40–42^

**Figure 4.**
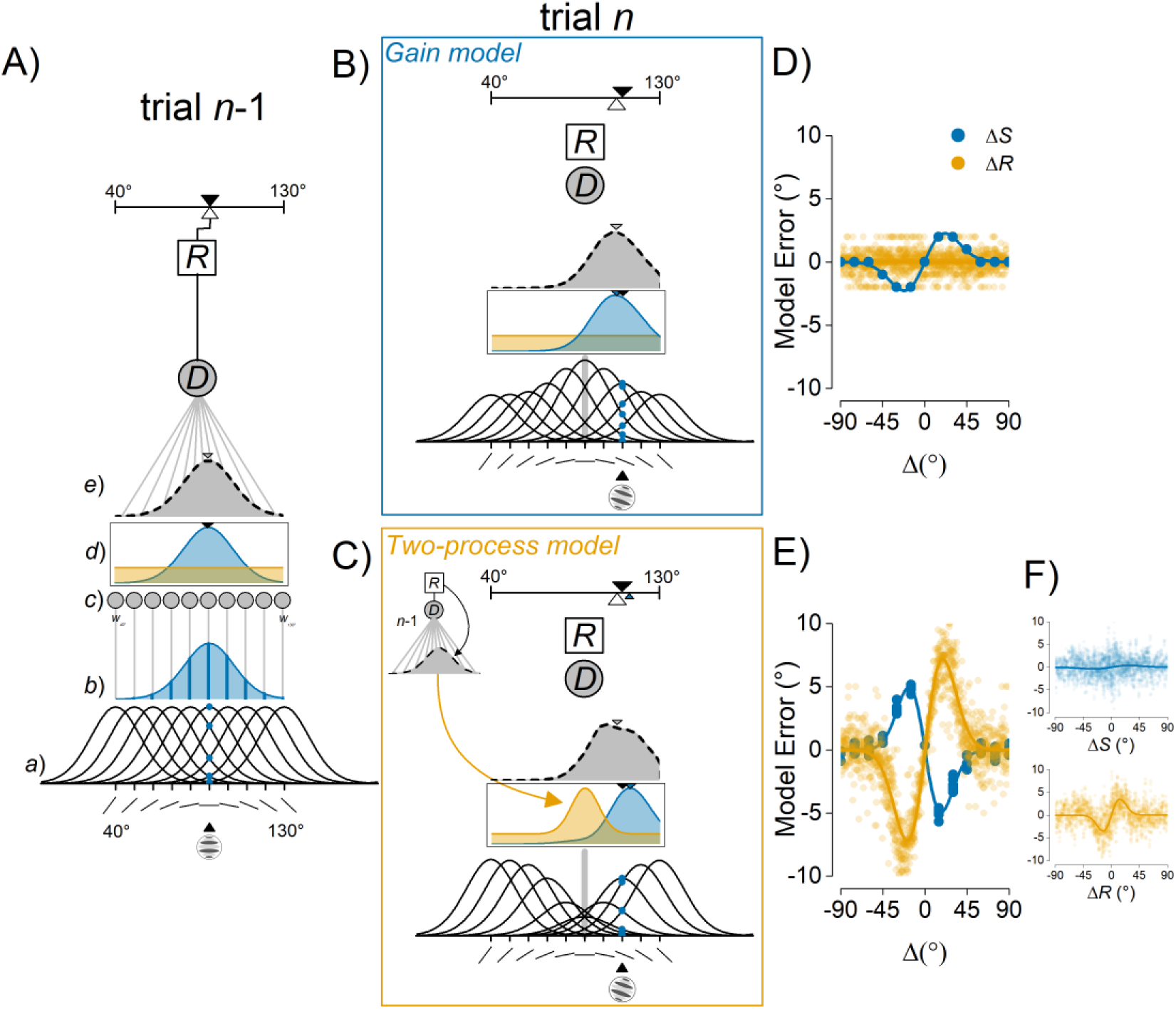
Predictions from two data-generating models of serial dependence. A) The general model architecture and the response to a first stimulus of 90° (trial n-1). From bottom to top: a) A subset of low-level orientation-selective units (from 40 to 130° in steps of 10°) with individual tuning curves around their preferred orientations and their individual response to the input stimulus (blue dots); *b*) Population response profile (blue distribution) with individual responses highlighted in dark blue; c) Connection weights (gray filled circles and lines) for the reading out of low-level activity by the decision unit; *d*) Population response (blue distribution) and uniformly distributed decisional weights (orange distribution); *e*) Weighted population response profile (gray distribution) decoded by the decision unit (*D*) and integrated into a final response (*R*). In the absence of any stimulus and response history, the reported orientation matches the input stimulus (white open triangle and black triangle, respectively). B) The Gain model tested on trial *n* with an input orientation of 110°. A gain change attracts the new stimulus toward the orientation presented before (gray line at 90°) altering the population response profile (blue distribution). Decisional weights are uniform and constant in this model (orange distribution). C) The Two-process model tested on trial *n* with an input orientation of 110°. The previous decoding process infiltrates the processing of new stimuli by altering the read-out weights of low-level activity (non-uniform orange distribution). This way decisional traces contrast the repulsive effect of low-level adaptation (in blue) leading to positive serial dependence. D) Predictions of the Gain model for the result of Experiment 4, with Δ*S* dissociated from Δ*R*. E) Predictions of the Two-process model for the results of Experiment 4. In panels A-B-C, black triangles represent the input orientation, blue triangles the orientation encoded by low-level units, gray-filled triangles the orientation decoded by the decision unit and white open triangles the final response. F) The free parameters of the Two-process model were optimized to approximate the results of Experiment 2 for both Δ*S* (top plot) and Δ*R* (bottom plot).

The model’s architecture included a layer of low-level units resembling a population of orientation-selective neurons (Figure 4A[*a*]), a higher-level decisional unit (Figure 4A[*e*]) and a final response stage. The presentation of an oriented stimulus elicits a population response (Figure 4A[*b*]) in which the contribution of each low-level unit depends on the distance between the stimulus and the unit’s preferred orientation. The decision unit then decodes and integrates the population profile into a final response (the reported orientation). The decoding stage occurs through a set of connection weights (Figure 4A[*c,d*]), mediating the read-out of the decision unit from each orientation-selective cell ^33^. Thus, orientation discrimination was modeled as a two-stage process: an initial response by a population of perceptual units and a later weighted decoding by a higher-level decisional unit.

In the Gain model (Figure 4B), serial dependence was produced by a gain change in the response properties of orientation-selective units ^19^ Exposure to a tilted stimulus increased the sensitivity of low-level units coding the same and nearby orientations and this, in turn, biased the population response to an incoming stimulus toward the orientation coded in the past. In this model, the decision unit decoded the population profile with uniform and constant weights; thus, serial dependence only emerged from trial-by-trial gain modulations at the level of orientation-selective neurons.

In the Two-process model (Figure 4C), serial dependence resulted from the interaction between low-level adaptation mechanisms and changes in decisional weights. Contrarily to the Gain model, exposure to oriented stimuli inhibited the response of low-level units centered onto the same and nearby orientations. This, in turn, biased the population response to an incoming stimulus away from the present orientation ^43,44^, resembling negative aftereffects observed after both brief and prolonged adaptation periods ^45–47^ The critical aspect of the Two-process model was the plasticity at the level of decisional weights. In this model, previous decoding strategies persisted, generating a non-uniform distribution of weights with maxima at/near the orientation decoded and reported in the recent past. This way, opposite carry-over effects were modeled at different stages of the processing stream with negative-aftereffects arising from low-level adaptation and attractive biases emerging from decisional traces. Because the reweighting process favored the read-out from orientation channels that were more informative in the recent past, this effect compensated for trial-by-trial adaptation at lower level, leading to the positive serial dependence observed after behavioral responses (Figure 4F). Crucially, the reweighting process was triggered by the requirement - and execution - of a behavioral response, and therefore, no weights update followed the mere exposure to stimuli.

The Two-process model provides a compelling account for the coexistence of both positive and negative dependencies in visual perception ^30,31^; however, its plausibility and performance had to be evaluated and tested against the Gain model. To this aim, we compared the predictions made by the two models with the true pattern of errors made by participants in a new experiment (Figure 4D, E). In this experiment (Experiment 4, Figure 1B), fifteen participants were presented with a sequence of Gabors in each single trial and were asked to report the orientation of the last stimulus after the sequence ended (see Methods). Because previous responses were always interleaved with a sequence of new stimuli before the next to-be-reported orientation, this paradigm offered an ideal situation in which previous stimuli and decisions were dissociated and both positive and negative serial dependence could emerge. To evaluate the predictions made by the two models in this experimental context, we ran a simulation of Experiment 4 (*n* = 50000) and we conditioned the error generated by both models on previous stimuli and responses.

As expected, the Gain model predicted positive serial dependence for stimuli but no influence of previous responses (Figure 4D). On the contrary, previous stimuli and perceptual decisions had opposite effects on the errors generated by the Two-process model: while adaptation at the low level caused repulsive biases, prior decisional weights attracted the representation of incoming stimuli (Figure 4E). In order to select which data-generating process provided a better approximation of the mechanisms underlying serial dependence, we compared the outcomes of both models with the actual data from Experiment 4. Quite remarkably, the behavioral results resembled almost exactly the predictions made by the Two-process model. A significant negative serial dependence was observed in response to previous stimuli (Δ*S*, α = -1.07°, *p* < 0.01, peak at 17.8°), whereas positive carry-over effects emerged from previous responses (Δ*R*, α = 4.02°, *p* < 0.001, peak at 21.3°), and the magnitude of the two dependency effects was significantly different (α(Δ*R*) vs. α(Δ*S*), *p* < 0.001; Figure 5).

**Figure 5.**
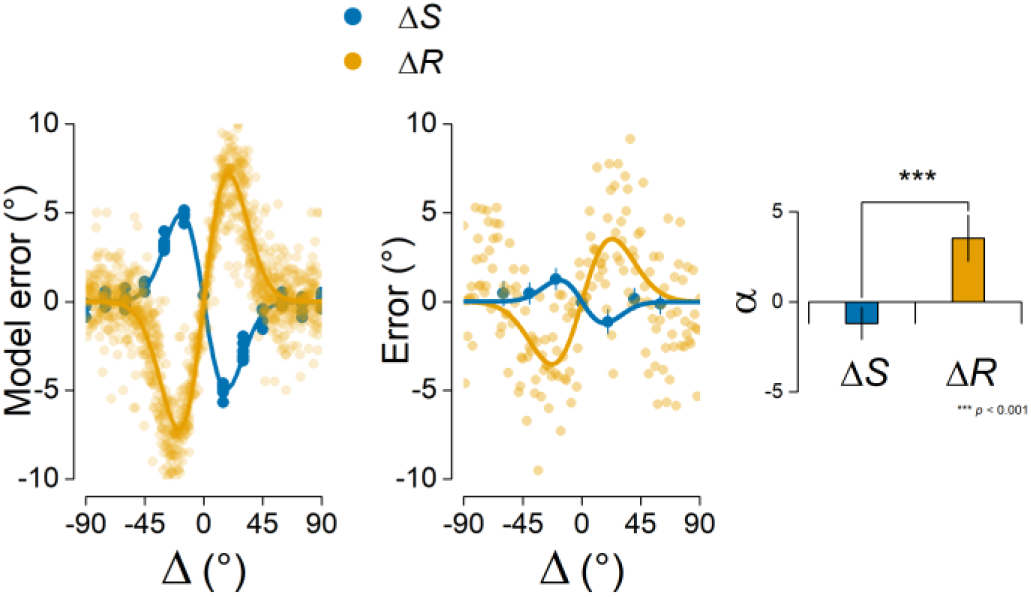
Opposite effects of previous stimuli and responses on perception. The observed results of Experiment 4 (middle and right panels) support the predictions of a Two-process model of serial dependence (left panel), with adaptation causing negative aftereffects (blue dots and line) and decisional traces inducing attractive biases (orange dots and line).

This clear-cut result provided strong evidence in favor of a hierarchical view of serial dependence and, in addition, it showed that decisional traces affect current perceptual decisions even when interleaved with new stimuli.

### Experiment 5: Location specificity of decisional traces

Having shown that attractive biases in perception emerge from the inertia of post-perceptual decoding mechanisms, in the present experiment (Experiment 5) we further characterized this phenomenon by investigating the specificity of decisional traces with regard to spatial location. Previous studies have shown stimulus-dependent carry-over effects that decreased with the spatial distance between consecutive stimuli, falling off for stimuli separated by 10-15° ^19,30^. Here we compared the spatial specificity of serial dependence when errors are conditioned on previous stimuli or previous responses.

Eighteen participants performed the orientation task with Gabors presented randomly at two peripheral locations (±6.5° from the center of the screen, along the horizontal meridian). With a cross-location spatial separation of 13°, serial dependence should significantly decrease when stimuli change their location across consecutive trials ^19^ However, if the spatial tuning of decisional traces differs from that measured for physical stimuli, a different pattern should emerge when errors are conditioned on previous responses.

The magnitude of serial dependence for both Δ*S* and Δ*R* was contrasted in two conditions, depending on whether two consecutive stimuli (with the ensuing responses) occurred at the same or at a different location. When Δ*S* was considered, serial dependence was significant for stimuli occurring at the same location (α = 2.13°, *p* < 0.01, peak at 30.9°) but absent for stimuli presented 13° apart (α = 0.71°, *p* = 0.10), thus confirming a significant effect of spatial distance (same vs. different location, *p* < 0.05; Figure 6A). However, when errors were conditioned on ΔR, serial dependence was present for stimuli both at the same (α = 2.58°, *p* < 0.001, peak at 21.5°) and at a different location (α = 4.05°, *p* < 0.001, peak at 21.5°). Moreover, despite the magnitude of the effect was numerically higher in the same-location condition, there was no reliable difference between the two (*p* = 0.09; Figure 6A). Thus, decisional traces have a location-independent tuning function that extends beyond that measured for previous stimuli.

**Figure 6.**
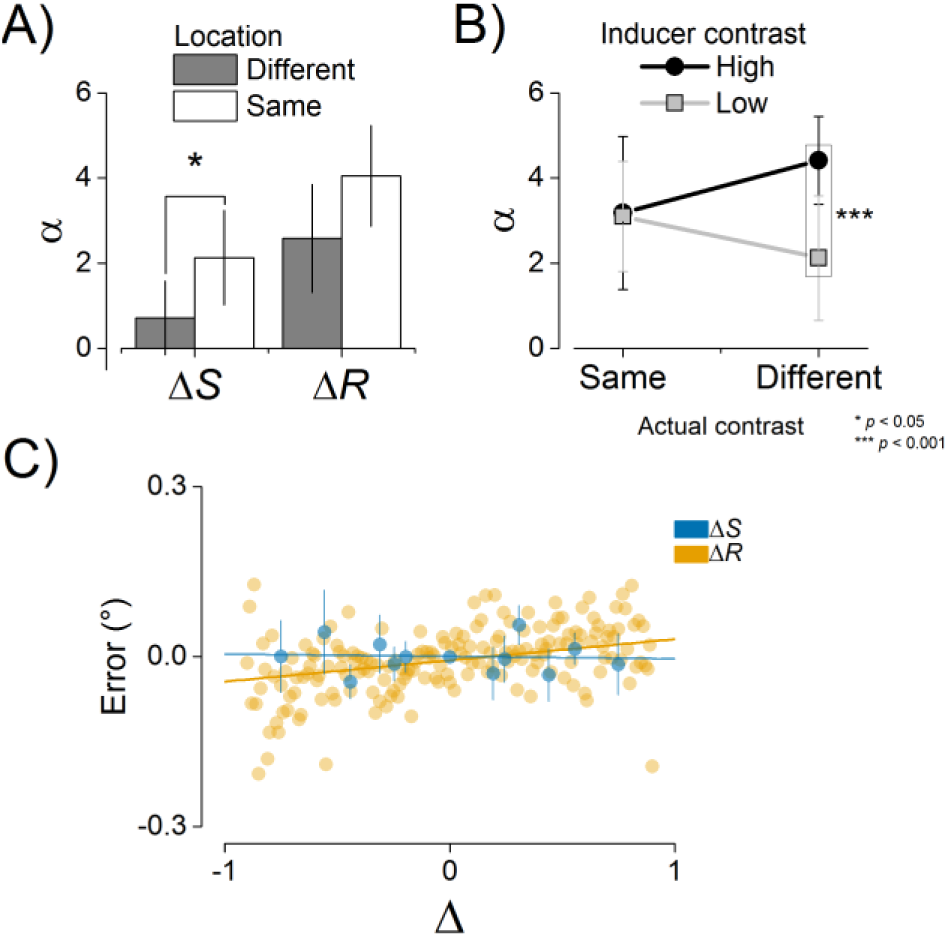
Characterization and generalization of decisional traces. A) Results of Experiment 5: Serial dependence disappeared for stimuli presented 13° apart, whereas previous responses significantly affected perception independently of the spatial location. B) Results of Experiment 6: High contrast inducer stimuli increased the impact of previous decisions on the processing of low-contrast stimuli. C) The effect of ΔS and ΔR in the ensemble coding task of Experiment 7: judgments of the average size were consistently biased by previous responses (Δ*R*, orange dots and line), whereas there was no influence of the average size of past ensembles (Δ*S*, blue dots and lines). Lines are best-fitting regression lines from multilevel linear modeling. Blue bars are 95% confidence intervals of the mean.

### Experiment 6: Stimulus strength and decisional traces

The fact that the lingering effect of previous responses persisted across spatial positions and time, even after streaks of intervening stimuli, is in line with the idea that positive serial dependence emerges from the inertia of a high-level decision unit, which operates in a location- and stimulus-independent manner. An interesting question is therefore whether decisional traces, once formed, become mere abstractions of a - biased - sensory experience or still preserve some low-level properties of the sensory evidence upon which they are based.

In the present experiment (Experiment 6) we measured serial dependence from previous responses as a function of stimulus contrast. The aim was two-fold: 1) we evaluated the impact of previous responses as a function of the actual sensory evidence, as determined by the stimulus visibility; and 2) we addressed whether decisional traces only rely on perceptual reports (the orientations confirmed with the response bar) or their impact varies with the strength of the previously decoded stimulus.

Fifteen participants performed the same orientation adjustment task as in Experiment 1, with stimuli of high (75%) and low (25%) contrast randomly intermixed across trials. In a first multilevel model, we compared the effect of switching between low and high stimulus contrast by extracting the α parameter when a low contrast inducer was followed by high contrast stimuli, and vice versa when a high contrast inducer was followed by low contrast stimuli. This analysis revealed significant serial dependence for ΔR in both cases (low-to-high, α = 1.97°, *p* < 0.01, peak at 24°; high-to-low, α = 4.16°, *p* < 0.001, peak at 24°) and a significant difference between the two conditions (*p* < 0.01). To further characterize this interaction between the contrast of previous and present stimuli, we then estimated the magnitude of serial dependence at the single-subject level, in order to compare four conditions of interest in a full factorial design. Individual α were submitted to twoway repeated-measures ANOVA with factors Inducer Contrast (Low vs. High), Actual Contrast (Same (as the inducer) vs. Different) and their interaction. The ANOVA yielded a significant main effect of the Inducer Contrast (F(_1,14_) = 6.11, *p* < 0.05, *η*_p_^2^, = 0.304) and a significant interaction Inducer Contrast x Actual Contrast (F(_1_,_14_) = 4.61, *p* < 0.05, *η*_p_^2^, = 0.248) driven by the significant difference between high-to-low and low-to-high contrast serial dependence (*p* < 0.001, Bonferroni correction; Figure 6B). Thus, decisional traces are stronger after high contrast stimuli and their attractive influence increases in relation to weak stimuli.

### Experiment 7: Generalization of serial dependence to ensemble coding

To generalize our findings to domains other than orientation perception, we performed a further experiment (Experiment 7) in which the task required participants to reproduce the average size of an ensemble of circles (see Methods and Figure 1C). Perceptual averaging, also defined as statistical representation or ensemble coding ^48,49^, refers to the ability of our perceptual system to extract global statistics, such as the mean or variability, from sets of similar stimuli, thus providing a coarse gist of complex visual scenes. It has been recently shown that ensemble coding occurs in space as well as in time and, consequently, the perceived statistics of an ensemble depends on its global properties in the immediate past ^50^. These results were discussed as evidence of a spatio-temporal continuity field in visual perception that promotes objects stability and coherence over time. However, based on the evidence reported thus far, it is plausible that serial dependence in summary statistic is due to the same post-perceptual mechanism that we have characterized for orientation processing.

To test this hypothesis, we predicted errors in the reported average size of fifteen new participants with two different multilevel linear models. In one model (Δ*S*), we used the difference between the average size of the previous and present ensemble as the dependent variable. In the other model (Δ*R*) we used the difference between the reported average size in the last trial and the current average size of the ensemble. The Δ*R* model was superior to the Δ*S* model in the overall fit (Akaike Information Criterion (AIC) for Δ*S* = 2000; AIC for Δ*R* = 1964) and previous responses were significantly (β = 0.037, *p* < 0.001) associated with errors of reported size. On the contrary, in the Δ*S* model the effect of previous stimuli were not significant (*p* > 0.6), indicating no systematic influence of previous ensembles on the reported average size (see Figure 6C).

## Discussion

The content of visual perception is strongly permeated by information lingering from the past. In the present study, we demonstrated that vision is shaped by two contrasting forces, arising from the history of physical stimuli and perceptual reports. Our work, based on multilevel modelling and cross-validation approaches, provides a thorough and comprehensive characterization of visual serial dependence from which we derived the following conclusions: 1) Visual perception is biased away from *previous stimuli*, 2) *Perceptual decisions* are serially dependent and 3) These two phenomena coexist and interact in shaping our interpretation of the visual input.

Our primary aim was to compare the ability of previous stimuli and responses to predict participants’ errors in orientation adjustment tasks. This comparison was performed in the first two experiments and showed, without ambiguity, that serial dependence in visual perception is explained by the history of behavioral responses and perceptual decisions. Previous responses outperformed past stimuli in predicting participants’ errors for both peripheral and foveal stimuli. In the latter case, the influence of previous stimuli was virtually absent. This indicated that response-induced history biases are independent of stimulus uncertainty and persist even when the perceptual resolution of stimuli increases and stimulus carry-over effects vanish.

In a following step, we hypothesized that serial dependence occurs exclusively after stimuli pass through a complete post-perceptual and decisional processing. Interestingly, the results of Experiment 3 revealed the opposite effects of stimulus and response history: After catch trials, in which perceptual decision was withheld because of task instructions, past stimuli evoked repulsive biases on subsequent orientation judgments; after response trials, however, adjustment errors were systematically attracted toward previous responses.

The repulsive impact of previous stimuli closely resembled the well-known tilt-aftereffect (TAE) observed after orientation adaptation ^28,46,46,51^. Previous work has shown that repulsive biases caused by the TAE are evident after few milliseconds of exposure to an oriented stimulus ^45,46,52^, suggesting that this rapid form of adaptation-induced plasticity may reflect an instantaneous and stereotyped behaviour of ensembles of sensory neurons. In line with this possibility, we demonstrated that previous stimuli, when isolated from the history of decisional processes, evoked systematic negative aftereffects ^30^.

Our crucial finding is that positive serial dependence required sequential decisions and perceptual reports. The most compelling evidence emerged from the results of Experiment 4, in which past stimuli and responses were dissociated and both positive and negative aftereffects could be assessed in a single perceptual report. Conditioning errors on past stimuli and responses in this key experiment showed the independent profile of the two sources of serial dependence, replicating the repulsive biases for stimuli observed in Experiment 4 and revealing the attractive impact of decisional traces even after streaks of intervening stimuli.

Thus, positive serial dependence is a post-perceptual phenomenon. A similar conclusion was reached by Fritsche, Mostert and DeLange^30^ using an interleaved procedure with adjustment and orientation-discrimination trials (two-alternative forced choice, 2AFC). The authors reported a task-dependent inversion of the effect of previous stimuli on current perception and decisions, with repulsion in 2AFC trials and attraction in adjustment responses. They concluded that positive serial dependence from past stimuli occurs at post-perceptual stages and may be mediated by biases in working memory. This implies that both attractive and repulsive history biases are induced by previous stimuli and their occurrence depends on the task at hand: when only perceptual processes are measured (e.g., in 2AFC), previous stimuli repel perception whereas when post-perceptual processes are tapped (e.g., using adjustment responses), previous stimuli attract decisions. However, this interpretation neglects the fact that even 2AFC tasks are strongly permeated by history biases arising from previous decisions/responses more than previous stimuli ^53^, and even more importantly, it overlooked the possibility that perceptual and post-perceptual processes may have their own, independent history-dependent biases.

Here we demonstrated the existence of two, functionally segregated processes of serial dependence which operate at different stages of the processing hierarchy. We show that attractive biases are not induced by recent stimuli on current responses but are the by-product of chains of decisional processes which are paralleled by repulsive adaptation at a lower stage.

To account for the opposite effects of stimuli and decisions, we proposed a Two-process model of serial dependence, combining low-level orientation adaptation and high-level decisional biases in a single hierarchical structure. We modelled low-level adaptation as a repulsive re-organization of a population of orientation-selective neurons ^43,54^ and the decisional stage as a weighted decoding of low-level information ^55–57^. By incorporating a memory trace for read-out weights at the decisional stage, we provided a simple model which approximated both low-level adaptation, in the absence of decisional activity, and attractive biases at post-perceptual stages. Crucially, the constant reweighting process counteracted and masked the effects of orientation adaptation, leading to the positive serial dependence reported here and in previous work ^19,30,34^ The Two-process model outperformed the Gain model of serial dependence proposed by Fischer and Whitney ^19^ in its ability to predict the outcomes of Experiment 4.

The plausibility of our Two-process model is corroborated by extensive evidence of read-out processes and integration of sensory information at high-level stages of processing ^55,58,59^ and by the critical role of decisional circuits in recent models of visual perception ^60,61^. Interestingly, read-out units, operating at later stages of the visual hierarchy, may have broader and largely location-independent tuning functions ^60,61^. This is in line with our results from Experiment 5, where we observed a broad spatial tuning of the effect of previous responses, which extends beyond the one reported for previous stimuli ^19,30^.

Although the Two-process model represents a parsimonious and unified account of serial dependence, other alternatives can be formulated. For example, by substituting decisional weights with probability distributions (priors) and low-level population activity with sensory evidence (likelihoods), one can accommodate our hierarchical model into the predictive coding framework ^62–65^. According to this perspective, higher-level probability distributions, built upon past information (e.g., previous stimuli), would be used to predict and test the incoming sensory input. As a consequence, when two consecutive stimuli are similar in some feature domain (e.g., orientation), the product of prior knowledge and current sensory evidence would result in mixed representations. It has been proposed that serial dependence emerges from such a predictive mechanism, educated by the fact that objects in the real world tend to remain stable over time. This implies that in the absence of a reliable temporal structure, the best prediction of future stimuli is that they will match the current percept^25^. Although this represents a suggestive theoretical proposal, it relies on the strong assumption that the brain systematically expects the reoccurrence of the same stimulus in a random sequence. However, if the past in a random series is used as a reference to build predictions, it seems more plausible that perception, shadowing the human reasoning system, would fall into the “local representativeness” heuristic ^66^, expecting consistent changes within short sequences of perceptual events ^8,67^ in order to match an internal model of randomness ^68,69^.

We argue that positive serial dependence may result from inertia in the reweighting process at decisional stages ^35^. According to the physics definition, we refer to “inertia” as the tendency of an object to resist changes in its actual state, which is proportional to its mass. This definition fits well with the results of our Experiment 6, demonstrating that decisional traces persist more strongly after the sensory/decision circuit has been perturbed with salient (high contrast) stimuli. One may therefore speculate that the stronger the intensity (mass) of a past perceptual episode, the larger its impact on future stimuli (inertia). However, this hypothesis is in sharp contrast with a recent report showing the opposite pattern in a 2AFC, that is, the tendency of choice repetitions to increase after weak sensory evidence ^35^ This study used different stimuli and, more importantly, a binary choice task which makes it difficult to compare to our results. One possibility is that decision boundaries and weights are differently distributed in tasks requiring a qualitative scaling (e.g., orientation adjustment) or comparative judgments, and this may alter the way decisions exert carry-over effects. This discrepancy may be an interesting question for future studies.

In a final experiment, we generalized our findings to the domain of ensemble coding ^49^, demonstrating that the appearance of an ensemble summary is biased toward past decisions while it is not affected by the statistical properties of previous stimuli ^50^.

What is the functional role of the two sources of serial dependence? We argue that both mechanisms serve the adaptive function of maximizing perceptual efficiency over the short-and long-term range. In free viewing conditions, we tend to make relatively rapid and large saccadic eye movements with subsequent fixations likely to land on stimuli with highly dissimilar structures ^45^. The rapid and repulsive reorganization of low-level sensory neurons may thus prepare our perceptual system to decorrelate and resolve different spatial structures, enhancing its ability to discriminate local features in a discontinuous perceptual environment. The lingering of decisional traces, on the other hand, may be explained by the integrative nature of decisional processes. Decisions terminate in a choice/response after sensory evidence is accumulated to some threshold level ^70^. When the action corresponding to the decision is postponed or the system is seeking for further evidence, the current state of the decisional process (i.e., the distribution of read-out weights) may be impressed into a temporary memory storage, which in turn becomes the starting point for collecting new evidence about future stimuli. Although the establishment of decisional weights is maladaptive in the short-term, leading to positive serial dependence, it may aid the long-term calibration of optimal decisional weights that fosters perceptual learning ^58,60,61^. Another intriguing possibility is that the decisional system implicitly counteracts adaptation and compensates for repulsive biases at the lower stage in the attempt to promote visual stability in a top-down fashion.

To conclude, in the present work we provide a unified account of the concatenated history of perceptual and decisional processes. We present a hierarchical model of serial dependence in which two separate and contrasting forces, exerted by adaptation and decision inertia, interact in shaping visual experience and our reactions to environmental stimuli. Building on recent work ^30^, our study has strong theoretical and experimental implications in dissociating different sources of history biases in perceptual and psychophysics research.

## Methods

### General Methods

A total of 91 subjects (62 females, mean age = 22.78±2.88), 55 from the University of Verona, Italy and 36 from the University of Fribourg, Switzerland, participated in this study for monetary payment (7€) or course credits. All participants had normal or corrected-to-normal vision and were naïve as to the purpose of the experiments. The study was carried out in accordance with the Declaration of Helsinki.

Stimuli were presented on a Multiscan G500 CRT 21” monitor (1024 × 768, 100 Hz, Experiment 1, 3, 5 and 6) and on a Philips 202P7 CRT (1600 × 1200, 85 Hz, Experiment 2, 4 and 6) and were generated with a set of custom-made programs written in Matlab and the Psychophysics Toolbox3.8^71^, running on Windows XP-based machines. All experiments were performed in dimly lit rooms and participants sat at a distance of 45 cm (Experiment 1, 3, 5 and 6) or 70 cm (Experiment 2,4 and 7) from the computer screen, with their head positioned on a chinrest. All stimuli were presented on a gray background (~24 cd/m^2^). Across experiments, the number of trials varied in order to achieve a minimum of 20 data points for each variable of interest, while keeping the overall duration within roughly one hour. Participants were instructed to maintain their gaze at the center of the screen for the entire duration of all experiments.

#### Experiment 1

An example of a trial sequence in Experiment 1 is illustrated in Figure 1A. Each trial started with a green central fixation spot (0.5°) followed by the presentation of a peripheral Gabor (8.5° to the left or right of fixation). Gabor stimuli had a peak contrast of 50% Michelson, spatial frequency of 0.5 cycles per degree and a Gaussian contrast envelope of 1.5°. After 400 ms, a noise mask appeared at the same location of the Gabor for 400 ms. Masks were white noise patches of the same size and peak contrast of the Gabor stimuli, smoothed with a symmetric Gaussian low-pass filter (filter size = 1°, s.d. = 2°). After a blank fixation interval of 500 ms, a response bar was presented at the same location of the Gabor. The response bar was a 0.5° wide white bar windowed in a Gaussian contrast envelope of 1.5°, with the same peak luminance of the Gabor stimuli. On each trial, the response bar appeared with a random orientation and participants were asked to rotate the bar until it matched the perceived orientation of the Gabor. The response bar was rotated by moving the computer mouse in the upward (clockwise rotation) or downward (counter-clockwise rotation) directions and the final response was confirmed by clicking the mouse left button. After a random interval (400-600 ms) a new trial started.

On each trial, the Gabor was assigned one of eighteen possible orientations, ranging from 0° (vertical) to 170° in steps of 10°. The series of orientation was pseudorandomly determined in order to keep the relative orientation between consecutive stimuli (Δ*S*, previous orientation minus present orientation) in the range ± 50°, in steps of 10°.

At the beginning of the experiment participants were provided written instructions and performed 20 trials of practice that were not recorded. The experiment consisted of four blocks of 140 trials each, for a total of 560 trials (~30 trials for each orientation tested and ~50 trials of each level of Δ*S*) and lasted approximately one hour. Stimuli were presented to the right or left of fixation in separate blocks. The average reaction time in all experiments using the same orientation adjustment task (Experiment 1-6) was 1.54 ± 0.42 seconds.

#### Experiment 2

The procedure in Experiment 2 was similar to Experiment 1 with the following exceptions. Gabor stimuli were presented at the center of the screen and had higher spatial frequency (1.2 cycles per degree). The orientation of stimuli varied pseudo-randomly in the range 0-179° (steps of 1°) and the relative orientation Δ*S* was a continuous variable ranging from -80° to +80° in steps of 1°. There were four blocks of 110 trials for a total of 440 trials.

#### Experiment 3

The procedure in Experiment 3 was similar to Experiment 1 with the following exceptions. In order to compare our results with previous work showing serial dependence in the absence of prior motor responses ^19^, we made our paradigm more similar to the one adopted by Fischer and Whitney (2014, Experiment 2). Gabor stimuli were presented at ± 6.5° from fixation for 500 ms and the duration of the mask was increased to 1000 ms. In about 40% of the total trials (catch trials), the response bar was replaced by a black disk (1.5° of diameter, 200 ms) explicitly instructing participants to not report the Gabor orientation. The duration of inter-trial intervals in catch trials was a running average of individual reaction times during the task. The orientation tested ranged from 0° to 165° in steps of 15°, with Δ*S* of ± 60° in steps of 15°. The increase of the step size for orientations and Δ*S* with respect to Experiment 1 allowed us to collect a sufficient number of data points for response trials (~30 for each orientation and ~40 for each Δ*S*) and catch trials (~20 for each orientation and ~26 for each Δ*S*) while keeping the overall duration of the experiment at about one hour.

#### Experiment 4

To test the predictions made by the two data-generating models of serial dependence, we performed an experiment in which stimuli to be reported were interleaved with sequences of stimuli not requiring a behavioral response. In this experiment we pursued an additional aim to measure the role of expectations in serial dependence comparing response biases after regular (i.e., ABCABC) or random orientation sequences. However, we found no differences between conditions and no explicit learning of the regular sequences, indicating that the employed manipulation of expectation was not effective. Therefore, given the suitability of this paradigm for evaluating the Two-process hypothesis of serial dependence, we used all data from this experiment to test the predictions made by the two data-generating processes.

An example of a trial sequence in Experiment 4 is illustrated in Figure 1B. Gabor stimuli were the same as in Experiment 2 and were presented at the same central location. Eighty percent of trials contained a sequence of six Gabors (200 ms each) with varying orientations (from 0° to 160° in steps of 20°) and a Δ*S* of ± 80°, in steps of 20°. No noise mask was presented after each stimulus and the phase of the sinusoidal component of each Gabor was changed randomly on each presentation (0-360°, steps of 15°). On the remaining 20% of total trials (catch trials) the sequence ended at a random position before the sixth Gabor (after 1-5 Gabors). These catch trials were used to control that participants were paying attention to the entire sequence and, most importantly, to the fifth stimulus before the to-be-reported orientation (5^th^ vs. 6^th^ stimulus error magnitude, *t*(_1,14_) = 0.21, *p* = 0.83, two-tailed t-test). After 400 ms from the offset of the last Gabor in the sequence, the response bar appeared at the center of the screen and participants had to report the orientation only of the most recently seen stimulus. Only trials where the sequence completed after six Gabors were analyzed.

There were four blocks of 71 trials each, for a total of 284 trials.

#### Experiment 5

The procedure, orientation steps and Δ*S* were the same as in Experiment 3. Gabor stimuli (1° of Gaussian envelope, spatial frequencies of 1 cycle/degree) were presented at ± 6.5° from fixation (as in Experiment 3) but their location (left or right) was randomly determined on each trial. Each Gabor lasted 300 ms and was followed by a noise mask for 500 ms.

The experiment consisted of a unique block of 240 trials, lasting approximately 20 minutes.

#### Experiment 6

The procedure, stimuli, orientation steps and Δ*S* were the same as in Experiment 3. The contrast of each Gabor was varied randomly from low (25% Michelson) to high (75% Michelson) on a trial-by-trial basis. The duration of the Gabor was also increased to 1000 ms and the following noise mask lasted 500 ms.

There were 5 blocks of 80 trials in Experiment 6, for a total of 400 trials (~ 1 hr).

#### Experiment 7

An example of a trial sequence in Experiment 7 is illustrated in Figure 1C. Stimuli were ensembles of sixteen light-gray dots of four different sizes. The average diameter of the ensemble was 0.75°, 0.945°, 1.19° or 1.5° and was randomly assigned on each trial (see Ariely, 2001 for a similar procedure). The standard deviation of the dots diameter within an ensemble was either 0.2° or 0.4°. The dots were arranged in a 4x4 matrix (14.1x10°), separated from the monitor’s center by 1.5° in the horizontal direction and 2° in the vertical direction. Dots were centered on equally spaced cells of 3.52x2.5°, with a random horizontal and vertical jitter ranging from 1 to 16 pixels. The distance of the ensemble’s center from the fixation spot was 11.14°.

Each trial started with a green fixation spot (0.5°) at the center of the screen for 400 ms. Then the ensemble appeared for 200 ms at one of four possible locations (top left, top right, bottom left, bottom right). After the ensemble offset, a dark-gray response dot appeared at the center of the screen with a random diameter (ranging from 0.2° to 4°) and participants were asked to adjust the diameter of the response dot to the perceived average size of the ensemble. The adjustment response was performed by moving the computer mouse in the upward (diameter increase) or downward (diameter decrease) directions and then clicking the left button to confirm the response. After a variable interval (500-700 ms) a new trial started. The average reaction time of participants in this experiment was 1.91 ± 0.52 ms.

The experiment consisted of 4 blocks of 120 trials, for a total of 480 trials (~50 minutes).

### Data Analysis

#### Outlier correction

As a first step in the analysis of the data from all experiments, circular errors in the orientation tasks (Experiment 1-6) were bounded to ±90° with respect to the stimulus orientation, approximating a normal distribution. Then, a Grubbs test^72^ was applied to exclude outliers from the adjustment errors of each participant (α = 0.05). For one subject in Experiment 3 we also excluded 4.5% of trials because of unrealistically fast adjustments (<0.5 sec). Finally, errors were demeaned to remove any chronic bias from the performance of each participant (e.g., systematic clockwise or counterclockwise errors in the orientation task or tendencies to report the diameter as larger or smaller in the ensemble coding task). As a consequence of this preprocessing, less than 2% of trials were excluded in total and the participants’ average absolute error was 8.04° ± 1.85 in the orientation tasks and 0.21° ± 0.03 in the ensemble coding task.

#### Multilevel Modeling

In all experiments, two competing models - corresponding to two different theoretical approaches – were built and compared. In the first model, errors were analyzed as a function of the difference between the orientation/average size of previous and present stimuli (Δ*S*). In the second model, the errors were analyzed as a function of the difference between previous responses and present stimuli (Δ*R*). The two predictors – i.e., the past response and its associated stimulus – were highly collinear and therefore it was not advisable to include them in the same model. Therefore, we fitted separate Δ*S* and Δ*R* models to all datasets and we compared goodness of their fit and the estimated parameters for all the experiments. To avoid the pitfalls of aggregated group-level data ^73–75^, all analyses (except one in Experiment 6) consisted of mixed-effects models with subjects as random effect ^76^.

#### Modeling orientation adjustment errors

In the orientation adjustment tasks of Experiment 1-2, serial dependence of the errors *y_ij_* was modeled with a nonlinear mixed-effects model of the form:

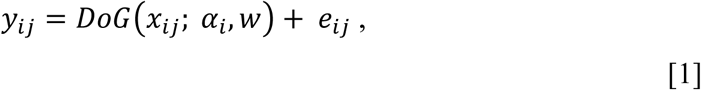

with *i* the subject index and *j* the *jth* trial and *DoG*(*x_ij_*; *α*, *w*) is given by:

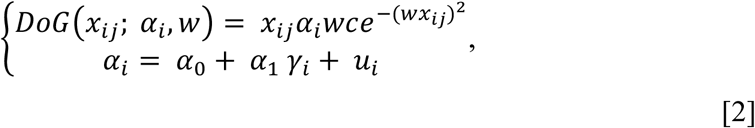

where *x_ij_* is the predictor variable (either Δ*S* or Δ*R*), (*α_i_*, *w*) are unknown parameters with *α_i_*∈ℝ the height of the peak and valley of the curve, *w* ∈ (0,1) the spread of the curve and 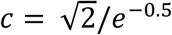 the normalization constant ^19^.*α_i_* is formed of a group-level (*α*_0_) component and an individual-level (*α*_1_) component with *γ_i_* being the subject indicator variable. *u_i_* is the group-level error assumed to be independent and identically distributed as 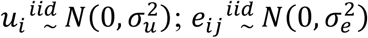 is the residual model error, independent of *u_i_*. The mixed-effects model analysis was performed using the *nlmefit* routine (*MaxIter* = 200, *TolX* = 1e^−4^, starting parameters: *α* = 2°, *w* = 0.05) and the *nlparci* function (for confidence intervals on parameters estimates) available in the MATLAB Statistics Toolbox ^77^.

Before model fitting, the random structure *u_i_* of the model was determined through a model selection procedure. Because of prior evidence of the effect of Δ*S* and the lack of investigation of the effect of Δ*R*, in the model selection for the random components we used Δ*S* as the independent variable *x*. Model selection was performed by comparing three models with all combinations of random effects. In the full model, both *α* and *w* were included as random parameters; in the two reduced models, *α* and *w* were included separately. The best random structure of the model was selected as the one with the lowest Akaike Information Criterion (AIC) in both Experiment 1 and 2. This procedure favored the model with only *α* as a random effect in both Experiment 1 (AIC(*α*) = 4.0653e+04; AIC(*w*) = 4.0684e+04; AIC(*α*+*w*) = 4.0655e+04) and Experiment 2 (AIC(*α*) = 2.0666e+04; AIC(*w*) = 2.0671e+04; AIC(*α*+*w*) = 2.0668e+04). Therefore, in all experiments and for both the estimation of Δ*S* and *ΔR* models we included only *α* as random effect.

In an effort to determine whether the serial dependence of the errors was associated with the physical stimuli presented in previous trials or with past perceptual decisions (i.e., responses in previous trials), in Experiment 1 and 2 we focused on an exhaustive comparison of the two competing models built with the Δ*S* and Δ*R* predictors. We proved which predictor provided a better fit of the data by means of Root Mean Squared error (*rms*). A non-parametric bootstrap test was used to determine whether there was a significant difference in the *rms* produced by the two Models (rms(AS-AR)). We first performed a Monte Carlo statistic by drawing 1000 bootstrap replicates of two null models (Δ*N*_1_, Δ*N*_2_), where the Δ*S* and Δ*R* predictors were randomly intermixed across trials: this way we obtained a surrogate null distribution of *rms(ANi-AN_2_)* in which differences in *rms* where not due to the separate ability of the Δ*S* and Δ*R* predictors to fit the data (see Figure 2A,B, top 78 histograms). A corrected p-value was then computed as the proportion of *rms*(Δ*N*_1_- Δ*N*_2_) that were higher or lower than the observed *rms*(Δ*S*-Δ*R*).

In a second step, to establish generalizability of the results, a cross-validation 79—82 approach was undertaken _79–82_. At each iteration of the process (n = 1000), the cross-validation procedure was based on four sequential steps: (1) The entire data set was randomly split into train (70% of the data) and test sets (30% of the data). The split was random but took into account the subject stratification (each subject contributed with an equal number of trials in the two sets); (2) The Δ*S* and Δ*R* models were separately fitted to the training data; (3) The parameters obtained in step 2 were then employed to generate predictions for the unseen data contained in the test set and (4) The *rms*(Δ*S*-Δ*R*) was calculated comparing the predicted vs. observed outcomes in the test set (see Figure 2A,B, bottom histograms). To obtain a p-value for the difference in the prediction accuracy between models, we calculated 1-*p_cv_* as the proportion of iterations in the cross-validation process in which the *rms*(Δ*S*-Δ*R*) was higher (*p_cv_*>0 = the Δ*R* model has lower *rms* and is favoured) or lower (*p_cv_*<0 = the Δ*S* model is favoured) than zero. This procedure gave a realistic estimate of how reliable is the superiority of the model based on Δ*R*.

The quantification of serial dependence for errors conditioned on both Δ*S* and Δ*R* was based on the estimated parameter *α*, in line with previous studies ^19,30,34^. Across all estimated models, the fixed-effect parameter *w* was 0.029 ± 0.011 for models with Δ*S* and 0.036 ± 0.006 for models with Δ*R*. The statistical significance of the *α* parameter was evaluated by calculating the z-score (the *α* coefficient divided by >83 its standard error) and converting it to a p-value using the standard normal curve ^83^.

To model serial dependence in experiments with two conditions (Experiment 3, 5 and 6) we adapted equation [1] to the following form:

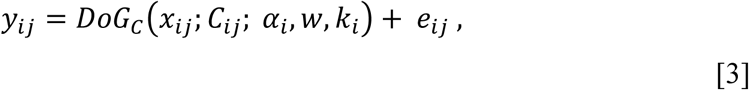

with the multilevel conditional derivative of Gaussian function *DoG_c_*(*x_ij_*; *C_ij_*; *α*, *w*, *k_i_*) given by:

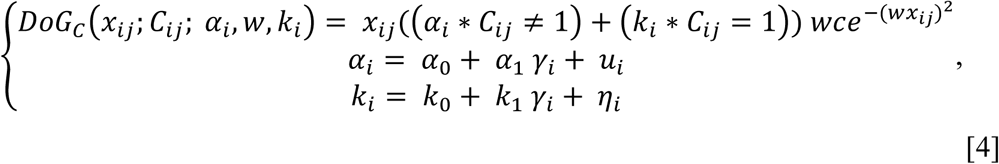

where *C_ij_* is a dummy variable [0,1] coding for the experimental condition (e.g., *C* = 0 for the Same-Location condition and *C* = 1 for the Different-Location condition of Experiment 5) and (*α_i_*, *k_i_*) are the independent height parameters of the curve in the two conditions (e.g., is the height for *C* = 0 and *k_i_* is the height for *C* = 1). As explained above, the height parameter(s) *(α_i_*,*k_i_*) have a fixed-effect component (*α*_0_, *k*_0_) that indicates group-level performances and an individual-level component (*α*_1_, *k*_1_) that indicates the deviation from the group-level performances. As per above the variable *γ_i_* is the individual-level indicator and *u_i_* and *η_i_* independent white noise processes. We also evaluated whether the inclusion of an additional parameter *z*, accounting for variations in the spread of the curve (*w*) as a function of *C* could improve the model fit. This was the case only for Experiment 3, where the AIC index was lower if *z* was included in the model (AIC *(α, w, k)* = 3.1085e+04; AIC(*α,w,k,z*) = 3.1080e+04), and therefore in this experiment we modeled the spread of serial dependence separately after Response-trials (*w* = 0.035) and after Catch-trials (*z* = 0.017). For the other experiments, there was a single fixed-effect *w*, while both the *α* and *k* height parameters were included as fixed and random effects, in line with the model selection above.

Statistical differences between the *α* and *k* parameters (differences between the magnitude of serial dependence across conditions) were assessed by computing a Z-Test of the estimated coefficients

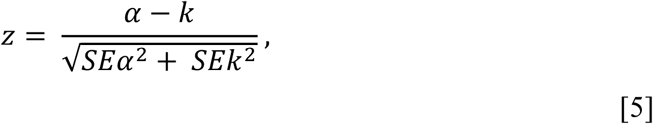

with *SEα* and *SEk* the standard error of the two parameters, and converting it to a p-value using the standard normal curve ^83^.

#### Orientation bias confound

It is well established that orientation judgments are subject to systematic biases in the form of repulsive shifts of the perceived orientation away from cardinal and oblique axes ^42,84^. Previous work has put forward the possibility that such biases, which are determined exclusively by the present stimulus orientation, may represent a critical confound in estimating serial dependence conditioned on previous responses ^85^. This bias takes a sinusoidal form over the orientation range and has been reported to produce spurious serial dependency effects on Δ*R*, when non-uniform stimulus orientations are presented in the trial sequence ^85^. Despite in our series of experiments the orientations were sampled from a uniform distribution covering the full 0-180° range, we wanted to further support the reliability of our findings by examining and ruling out this potential confound. To this aim, for all experiments with the orientation adjustment task we run a Monte Carlo bootstrap procedure in which we estimated the nonlinear mixed-effects model with Δ*R* 10000 times, shuffling the temporal order of the orientations (along with their associated responses) across trials. This way any temporal relationship (i.e., true serial dependence) between previous responses and actual stimuli was removed and we were able to estimate the spurious effects caused by the orientation bias alone, and to compare it with the magnitude of the α parameter observed when the temporal structure of the task was maintained.

Across all experiments we found a small spurious effect reflecting the orientation bias which led to an average magnitude of serial dependence of α = 1.48 ± 0.88°; crucially, however, the α values obtained from the shuffling procedure in all experiments were significantly lower than the observed ones (all *p* < 0.01, except in Experiment 2: *p* = 0.07; p-value obtained as the proportion of bootstrapped α higher than the observed one). This validation procedure, along with the generalization of the Δ*R* effect to a different task, free from orientation biases (Experiment 7), gives us strong confidence in the reported findings.

#### Modeling errors in ensemble coding

In the ensemble coding task of Experiment 7, the error *y_ij_* was modeled with a linear mixed-effects model of the form:

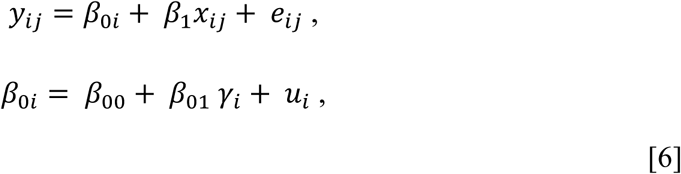

where *β*_1_ *and β*_0*i*_ are the slope and intercept and *x_ij_* the independent variable. The intercept *β*_0*i*_ had a multilevel structure with: (a) *β*_00_ representing the group-level fixed effect, (b) *β*_01_ representing the individual-level parameter which is coded in (c) the variable *γ_i_* and (d) an associated white noise parameter 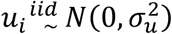. 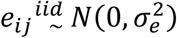 denotes the residual model error. The Δ*S* and Δ*R* predictors were included as the independent variable *x* in two separate models.

#### Data-generating processes of serial dependence

In the following we describe and evaluate the predictions of two opposite explanatory processes for serial dependence: The Gain model and the Two-process model. Both data-generating processes were composed by a three-layers architecture including: (1) a labeled line-like population of orientation selective units ^54^, (2) one decisional unit and (3) one response stage. Low-level units were 180 circular normal distributions centered on orientations from 0° to 180° in steps of 1° (Figure 4A[*a*]). The response of each unit, *r_θ_* to an input orientation *θ* was defined as:

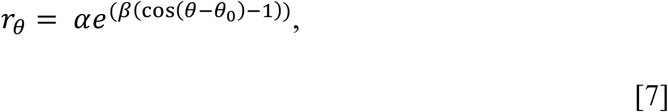

where *α* is the amplitude or peak response, *β* regulates the width of the tuning function and *θ*_0_ is the unit preferred orientation ^43,54^.

In the absence of any stimulus and response history, both models contemplated the same computation: all low-level units had parameter values of *α* = 1 and *β* = 4.68 (corresponding to a full width at half height of 28.2° ^19^). The collective low-level units’ response (population profile) was a vector of single-unit activations in which each channel signaled its preferred orientation with a value proportional to its response (the closer the input to the preferred orientation, the larger the response, see Figure 4A[*b*]). We modeled the decisional stage following previous work and assuming that a decision unit reads out and integrates the activity of a population of early sensory neurons ^55–57^. The decision unit in our model decoded low-level information by computing a weighted circular mean of the population profile ^54,60^ (see Figure 4A[*e*]), with baseline uniform weights of 0.0056 (integrating to 1 over the 180 orientations). The decoded orientation was then relayed to the response unit for the final outcome.

The Gain and Two-process model differed in their respective operating mechanisms once we start considering the temporal evolution of a series of stimuli and their associated responses. In the Gain model, a gain change was applied to the tuning function of low-level units after the presentation of each stimulus. The gain change was regulated by a circular distribution (eq. [7]) with *β* = 4.68 and *α_g_* (the gain factor) as a free parameter. This increased the responsivity of units centered on recently seen orientations (Figure 4B), producing the attractive bias of serial dependence ^19^ Decisional weights remained uniforms in the Gain model and had no effect on stimulus decoding.

In the Two-process model of serial dependence, low-level adaptation was regulated by the same circular distribution as in the Gain model but with inverted shape, inhibiting the response of units tuned to recently seen orientations. This caused the repulsive shifts in the responses of low-level orientation channels observed after adaptation ^54^ (see Figure 4C). The decoding (and report) of stimulus orientation altered the readout weights of the decision unit according to:

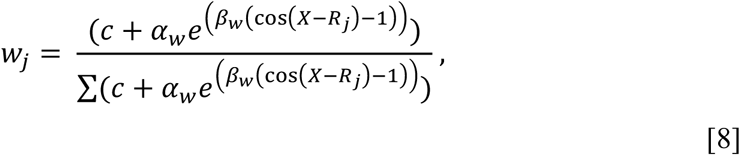

where the reweighting *w* in trial *j* is a function of *α_w_* and *β_w_*, two free parameters reflecting the peak and width of a circular distribution over the whole orientation range *X*, and the decoded/reported orientation *R_j_* which sets the center of the new weights distribution (see Figure 4C, yellow distribution). *c* constitutes a small baseline offset (*c* = 0.1) to avoid zero weights. The denominator is a normalization factor ensuring that the weights distribution integrates to 1.

The tenet of this model was that the reweighting process was triggered by behavioral responses and thus, occurred only for orientations requiring a perceptual report *R_j_*. Changes in decisional weights from trial to trial were then modeled with an update function:

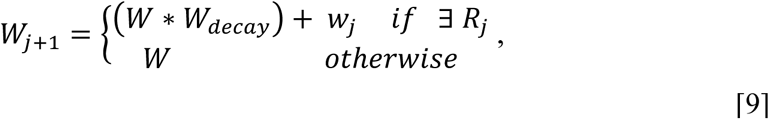

where *W*_*j*+1_ is the new set of weights affecting the readout of future stimuli, *W_decay_* is a free forgetting parameter, which determines how fast the history of weights *W* relaxes back to uniform and *W_j_* is the last reweighted distribution of eq. [8].

In an effort to evaluate the predictions of the two models, a simulation with 100000 trials was conducted. The models free parameters (*α_g_* [gain factor], *α_w_* and *β_w_*[peak and width of the weight distribution], and *W_decay_*[forgetting factor]) were estimated by means of least-squares minimization, in order to minimize the difference between the simulated amplitudes of serial dependence (*α*) for Δ*S* and Δ*R* and those observed in Experiment 2, where the paradigm (stimuli and foveal presentation) was similar to the one used in Experiment 4. A set of plausible parameters (*α_g_* [0.05-0.95, steps = 20], *α_w_* [0.05-0.95, steps = 20], *β_w_*[131:1.90, steps = 20], and *W_decay_*[0.05-0. 95, steps = 20]) was tested following a full factorial design (20x20x20x20). To approximate the results of Experiment 2, however, it was necessary to reproduce the different amplitude of serial dependence observed for Δ*S* (α = 0.30°) and Δ*R* (α = 3.40°). This was done using the Two-process model in the optimization procedure and adding a noise component *ω* to the response *R_j_*, with 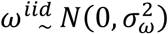 and *σ_ω_* = 20°, leading to an average absolute error of ~10°, close to the one observed across experiments (8.04°). Because the response on the last trial determined the center of the new weights distribution (eq. [8]), the additive noise *ω* randomly separated the reported from the input orientation, thus disentangling the response and the weights update from the stimulus. This additional noise component was only implemented in the parameters optimization based on Experiment 2, whereas it was not necessary to predict the results of Experiment 4 in which the responses and stimuli were already disentangled by the specific paradigm.

The optimization procedure led to a set of estimated parameters (*α_g_* [0.75], *α_w_* [0.425], *β_w_* [15.10], and *W_decay_* [0.05]) which approximated well the observed amplitudes of serial dependence in Experiment 2 (*α*(Δ*S*) = 0.32°, α(Δ*R*) = 3.23°; Figure 4F). These parameters were then used in both the Gain and the Two-process model to predict the results of Experiment 4 (see Results).

## References

1. Anstis, S., Verstraten, F. A. & Mather, G. The motion aftereffect. Trends Cogn. Sci. 2, 111–117 (1998).

2. Gibson, J. J. Adaptation with negative after-effect. Psychol. Rev. 44, 222 (1937).

3. Hayhoe, M. & Wenderoth, P. Adaptation Mechanisms in Color and Brightness. in From Pigments to Perception (eds. Valberg, A. & Lee, B. B.) 353–367 (Springer US, 1991). doi:10.1007/978-1-4615-3718-2_41

4. Gibson, J. J. & Radner, M. Adaptation, after-effect and contrast in the perception of tilted lines. I. Quantitative studies. J. Exp. Psychol. 20, 453 (1937).

5. Maloney, L. T., Dal Martello, M. F., Sahm, C. & Spillmann, L. Past trials influence perception of ambiguous motion quartets through pattern completion. Proc. Natl. Acad. Sci. U. S. A. 102, 3164–3169 (2005).

6. Kristjánsson, Á. & Campana, G. Where perception meets memory: A review of repetition priming in visual search tasks. Atten. Percept. Psychophys. 72, 5–18 (2010).

7. Maljkovic, V. & Nakayama, K. Priming of pop-out: I. Role of features. Mem. Cognit. 22, 657–672 (1994).

8. Pascucci, D., Mastropasqua, T. & Turatto, M. Permeability of priming of pop out to expectations. J. Vis. 12, 21–21 (2012).

9. Abrahamyan, A., Silva, L. L., Dakin, S. C., Carandini, M. & Gardner, J. L. Adaptable history biases in human perceptual decisions. Proc. Natl. Acad. Sci. 113, E3548–E3557 (2016).

10. Jesteadt, W., Luce, R. D. & Green, D. M. Sequential effects in judgments of loudness. J. Exp. Psychol. Hum. Percept. Perform. 3, 92 (1977).

11. Vinson, D. W., Dale, R. & Jones, M. N. Decision contamination in the wild: Sequential dependencies in Yelp review ratings. in Proceedings of the 38th Annual Meeting of the Cognitive Science Society 1433–1438 (2016).

12. Jonides, J. & Nee, D. E. Brain mechanisms of proactive interference in working memory. Neuroscience 139, 181–193 (2006).

13. Makovski, T. & Jiang, Y. V. Proactive interference from items previously stored in visual working memory. Mem. Cognit. 36, 43–52 (2008).

14. Rahnev, D., Koizumi, A., McCurdy, L. Y., D’Esposito, M. & Lau, H. Confidence leak in perceptual decision making. Psychol. Sci. 26, 1664–1680 (2015).

15. Wing, A. M. & Kristofferson, A. B. Response delays and the timing of discrete motor responses. Percept. Psychophys. 14, 5–12 (1973).

16. Levinson, E. & Sekuler, R. Adaptation alters perceived direction of motion. Vision Res. 16, 779–IN7 (1976).

17. Thompson, P. & Burr, D. Visual aftereffects. Curr. Biol. 19, R11–R14 (2009).

18. Kiyonaga, A., Scimeca, J. M., Bliss, D. P. & Whitney, D. Serial Dependence across Perception, Attention, and Memory. Trends Cogn. Sci. (2017).

19. Fischer, J. & Whitney, D. Serial dependence in visual perception. Nat. Neurosci. 17, 738–743 (2014).

20. Corbett, J. E., Fischer, J. & Whitney, D. Facilitating Stable Representations: Serial Dependence in Vision. PLOS ONE 6, e16701 (2011).

21. Cicchini, G. M., Anobile, G. & Burr, D. C. Compressive mapping of number to space reflects dynamic encoding mechanisms, not static logarithmic transform. Proc. Natl. Acad. Sci. 111, 7867–7872 (2014).

22. Liberman, A., Fischer, J. & Whitney, D. Serial dependence in the perception of faces. Curr. Biol. 24, 2569–2574 (2014).

23. Liberman, A. & Whitney, D. The serial dependence of perceived emotional expression. J. Vis. 15, 929–929 (2015).

24. Xia, Y., Leib, A. Y. & Whitney, D. Serial dependence in the perception of attractiveness. J. Vis. 16, 28–28 (2016).

25. Burr, D. & Cicchini, G. M. Vision: efficient adaptive coding. Curr. Biol. 24, R1096–R1098 (2014).

26. Dragoi, V., Sharma, J. & Sur, M. Adaptation-Induced Plasticity of Orientation Tuning in Adult Visual Cortex. Neuron 28, 287–298 (2000).

27. Huk, A. C., Ress, D. & Heeger, D. J. Neuronal Basis of the Motion Aftereffect Reconsidered. Neuron 32, 161–172 (2001).

28. Jin, D. Z., Dragoi, V., Sur, M. & Seung, H. S. Tilt Aftereffect and Adaptation-Induced Changes in Orientation Tuning in Visual Cortex. J. Neurophysiol. 94, 4038–4050 (2005).

29. Mather, G., Pavan, A., Campana, G. & Casco, C. The motion after-effect reloaded. Trends Cogn. Sci. 12, 481–487 (2008).

30. Fritsche, M., Mostert, P. & de Lange, F. P. Opposite effects of recent history on perception and decision. Curr. Biol. 27, 590–595 (2017).

31. Schwiedrzik, C. M. et al. Untangling perceptual memory: Hysteresis and adaptation map into separate cortical networks. Cereb. Cortex 24, 1152–1164 (2012).

32. Snyder, J. S., Schwiedrzik, C. M., Vitela, A. D. & Melloni, L. How previous experience shapes perception in different sensory modalities. Front. Hum. Neurosci. 9, (Front. Hum. Neurosci. 9,).

33. Dosher, B. A., Jeter, P., Liu, J. & Lu, Z.-L. An integrated reweighting theory of perceptual learning. Proc. Natl. Acad. Sci. 110, 13678–13683 (2013).

34. Liberman, A., Zhang, K. & Whitney, D. Serial dependence promotes object stability during occlusion. J. Vis. 16, 16–16 (2016).

35. Akaishi, R., Umeda, K., Nagase, A. & Sakai, K. Autonomous mechanism of internal choice estimate underlies decision inertia. Neuron 81, 195–206 (2014).

36. Mareschal, I. & Shapley, R. M. Effects of contrast and size on orientation discrimination. Vision Res. 44, 57–67 (2004).

37. Sally, S. L. & Gurnsey, R. Foveal and extra-foveal orientation discrimination. Exp. Brain Res. 183, 351–360 (2007).

38. Burr, D. C. & Wijesundra, S.-A. Orientation discrimination depends on spatial frequency. Vision Res. 31, 1449–1452 (1991).

39. Aiken, L. S., West, S. G. & Reno, R. R. Multiple regression: Testing and interpreting interactions. (Sage, 1991).

40. Ahissar, M. & Hochstein, S. The reverse hierarchy theory of visual perceptual learning. Trends Cogn. Sci. 8, 457–464 (2004).

41. Riesenhuber, M. & Poggio, T. Hierarchical models of object recognition in cortex. Nat. Neurosci. 2, 1019–1025 (1999).

42. Wei, X.-X. & Stocker, A. A. A Bayesian observer model constrained by efficient coding can explain’anti-Bayesian’percepts. Nat. Neurosci. 18, 1509–1517 (2015).

43. Clifford, C. W., Wenderoth, P. & Spehar, B. A functional angle on some after-effects in cortical vision. Proc. R. Soc. Lond. B Biol. Sci. 267, 1705–1710 (2000).

44. Pavan, A., Hocketstaller, J., Contillo, A. & Greenlee, M. W. Tilt aftereffect following adaptation to translational Glass patterns. Sci. Rep. 6, (2016).

45. Dragoi, V., Sharma, J., Miller, E. K. & Sur, M. Dynamics of neuronal sensitivity in visual cortex and local feature discrimination. Nat. Neurosci. 5, 883–891 (2002).

46. Dragoi, V. & Sur, M. Plasticity of orientation processing in adult visual cortex. Vis. Neurosci. 1654–1664 (2004).

47. Zavitz, E., Yu, H.-H., Rowe, E. G., Rosa, M. G. P. & Price, N. S. C. Rapid Adaptation Induces Persistent Biases in Population Codes for Visual Motion. J. Neurosci. 36, 4579–4590 (2016).

48. Ariely, D. Seeing sets: Representation by statistical properties. Psychol. Sci. 12, 157–162 (2001).

49. Haberman, J. & Whitney, D. Ensemble perception: Summarizing the scene and broadening the limits of visual processing. Percept. Conscious. Search. Anne Treisman 339–349 (2012).

50. Manassi, M., Liberman, A., Chaney, W. & Whitney, D. The perceived stability of scenes: serial dependence in ensemble representations. Sci. Rep. 7, 1971 (2017).

51. Pascucci, D. & Turatto, M. Immediate effect of internal reward on visual adaptation. Psychol. Sci. 24, 1317–1322 (2013).

52. Sekuler, R. & Littlejohn, J. Tilt aftereffect following very brief exposures. Vision Res. 14, 151–152 (1974).

53. Fründ, I., Wichmann, F. A. & Macke, J. H. Quantifying the effect of intertrial dependence on perceptual decisions. J. Vis. 14, 9–9 (2014).

54. Pouget, A., Zhang, K., Deneve, S. & Latham, P. E. Statistically efficient estimation using population coding. Neural Comput. 10, 373–401 (1998).

55. Gold, J. I. & Shadlen, M. N. The neural basis of decision making. Annu Rev Neurosci 30, 535–574 (2007).

56. Haefner, R. M., Gerwinn, S., Macke, J. H. & Bethge, M. Inferring decoding strategies from choice probabilities in the presence of correlated variability. Nat. Neurosci. 16, 235–242 (2013).

57. Nienborg, H. & Cumming, B. Correlations between the activity of sensory neurons and behavior: how much do they tell us about a neuron’s causality? Curr. Opin. Neurobiol. 20, 376–381 (2010).

58. Law, C.-T. & Gold, J. I. Reinforcement learning can account for associative and perceptual learning on a visual-decision task. Nat. Neurosci. 12, 655–663 (2009).

59. Roitman, J. D. & Shadlen, M. N. Response of neurons in the lateral intraparietal area during a combined visual discrimination reaction time task. J. Neurosci. Off. J. Soc. Neurosci. 22, 9475–9489 (2002).

60. Dosher, B. A., Jeter, P., Liu, J. & Lu, Z.-L. An integrated reweighting theory of perceptual learning. Proc. Natl. Acad. Sci. 110, 13678–13683 (2013).

61. Watanabe, T. & Sasaki, Y. Perceptual learning: toward a comprehensive theory. Annu. Rev. Psychol. 66, 197–221 (2015).

62. Kersten, D., Mamassian, P. & Yuille, A. Object perception as Bayesian inference. Annu Rev Psychol 55, 271–304 (2004).

63. Kok, P. & de Lange, F. P. Predictive coding in sensory cortex. in An introduction to model-based cognitive neuroscience 221–244 (Springer, 2015).

64. Rao, R. P. & Ballard, D. H. Predictive coding in the visual cortex: a functional interpretation of some extra-classical receptive-field effects. Nat. Neurosci. 2, (1999).

65. Summerfield, C. & De Lange, F. P. Expectation in perceptual decision making: neural and computational mechanisms. Nat. Rev. Neurosci. 15, 745 (2014).

66. Tversky, A. & Kahneman, D. Judgment under uncertainty: Heuristics and biases. in Utility, probability, and human decision making 141–162 (Springer, 1975).

67. Chopin, A. & Mamassian, P. Predictive properties of visual adaptation. Curr. Biol. 22, 622–626 (2012).

68. Croson, R. & Sundali, J. The gambler’s fallacy and the hot hand: Empirical data from casinos. J. Risk Uncertain. 30, 195–209 (2005).

69. Tversky, A. & Kahneman, D. Belief in the law of small numbers. Psychol. Bull. 76, 105 (1971).

70. Shadlen, M. N. & Kiani, R. Decision making as a window on cognition. Neuron 80, 791–806 (2013).

71. Brainard, D. H. & Vision, S. The psychophysics toolbox. Spat. Vis. 10, 433–436 (1997).

72. Grubbs, F. E. Procedures for detecting outlying observations in samples. Technometrics 11, 1–21 (1969).

73. Ashby, F. G., Maddox, W. T. & Lee, W. W. On the dangers of averaging across subjects when using multidimensional scaling or the similarity-choice model. Psychol. Sci. 5, 144–151 (1994).

74. Curran, T. & Hintzman, D. L. Violations of the independence assumption in process dissociation. J. Exp. Psychol. Learn. Mem. Cogn. 21, 531 (1995).

75. Estes, W. K. The problem of inference from curves based on group data. Psychol. Bull. 53, 134 (1956).

76. Baayen, R. H., Tweedie, F. J. & Schreuder, R. The subjects as a simple random effect fallacy: Subject variability and morphological family effects in the mental lexicon. Brain Lang. 81, 55–65 (2002).

77. Jones, B. MATLAB: Statistics Toolbox; User’s Guide. (MathWorks, 1997).

78. Davison, A. C. & Hinkley, D. V. Bootstrap methods and their application. 1, (Cambridge university press, 1997).

79. Browne, M. W. Cross-validation methods. J. Math. Psychol. 44, 108–132 (2000).

80. Hastie, T., Tibshirani, R. & Friedman, J. The Elements of Statistical Learning: Data Mining, Inference, and Prediction. Biometrics (2002).

81. Pitt, M. A. & Myung, I. J. When a good fit can be bad. Trends Cogn. Sci. 6, 421–425 (2002).

82. Stone, M. Cross-validatory choice and assessment of statistical predictions. J. R. Stat. Soc. Ser. B Methodol. 111–147 (1974).

83. Cohen, J., Cohen, P., West, S. G. & Aiken, L. S. Applied multiple regression/correlation analysis for the behavioral sciences. (Routledge, 2013).

84. Balikou, P. et al. Independent sources of anisotropy in visual orientation representation: a visual and a cognitive oblique effect. Exp. Brain Res. 233, 3097–3108 (2015).

85. Fritsche, M. & others. To Smooth or not to Smooth: Investigating the Role of Serial Dependence in Stabilising Visual Perception. From the 2 (2016).

